# Single-cell analysis reveals the function of lung progenitor cells in COVID-19 patients

**DOI:** 10.1101/2020.07.13.200188

**Authors:** Zixian Zhao, Yu Zhao, Yueqing Zhou, Xiaofan Wang, Ting Zhang, Wei Zuo

## Abstract

The high mortality of severe 2019 novel coronavirus disease (COVID-19) cases is mainly caused by acute respiratory distress syndrome (ARDS), which is characterized by increased permeability of the alveolar epithelial barriers, pulmonary edema and consequently inflammatory tissue damage. Some but not all patients showed full functional recovery after the devastating lung damage, and so far there is little knowledge about the lung repair process^1^. Here by analyzing the bronchoalveolar lavage fluid (BALF) of COVID-19 patients through single cell RNA-sequencing (scRNA-Seq), we found that in severe (or critical) cases, there is remarkable expansion of TM4SF1+ and KRT5+ lung progenitor cells. The two distinct populations of progenitor cells could play crucial roles in alveolar cell regeneration and epithelial barrier re-establishment, respectively. In order to understand the function of KRT5+ progenitors in vivo, we transplanted a single KRT5+ cell-derived cell population into damaged mouse lung. Time-course single-cell transcriptomic analysis showed that the transplanted KRT5+ progenitors could long-term engrafted into host lung and differentiate into HOPX+ OCLN+ alveolar barrier cell which restored the epithelial barrier and efficiently prevented inflammatory cell infiltration. Similar barrier cells were also identified in some COVID-19 patients with massive leukocyte infiltration. Altogether this work uncovered the mechanism that how various lung progenitor cells work in concert to prevent and replenish alveoli loss post severe SARS-CoV-2 infection.

---

COVID-19 caused by SARS-CoV-2 virus infection is the major health concern worldwide. Pathological studies of COVID-19 postmortem lungs have shown that the effect of mild virus infection is limited in upper airway and had little influence on the lung tissue integrity. However, severe virus infection leads to diffuse alveolar damage (DAD) characterized apoptosis, desquamation of alveolar epithelial cells, and infiltration of inflammatory cells into alveolar cavity, which could eventually lead to hypoxemia, pulmonary tissue fibrosis and death of patients. Hyperplasia of type II alveolar cells (ATII) was also noted in most cases, which could suggest an undergoing regenerative process mediated by ATII lung progenitor cells^2-4^. In order to fully elucidate the epithelial damage and repair mechanism, we analyzed the single cell transcriptomic profile of lung BALF to quantify the major events post infection and focused on structural epithelial cells. BALF is a useful technique for sampling the human lung, providing landscape information of the whole lower respiratory tract. The current study was based on public scRNA-Seq datasets on BALF cells from three patients with moderate COVID-19 (M1 – M3), six patients with severe/critical infection (S1–S6) and four healthy controls (HC1–HC4)^5,6^.

Firstly, we performed unsupervised clustering analysis on the whole dataset to separate EPCAM+/TPPP3+/KRT18+ epithelial cells from other cells types (mostly immune cells) in the BALF (Extended Data Fig.1a,b). Reclustering analysis identified 12 epithelial cell clusters, among them 4 were identified to be co-expressing immune markers which could be epithelial cells engulfed by leukocytes (Extended Data Fig. 1c,d). The other 8 distinct epithelial cell clusters composed of Club/goblet cells (Cluster 0. SCGB1A1+/MUC5AC+), various types of ciliated cells (Cluster 1-5. FOXJ1+), alveolar cells (Cluster 6. HOPX+/SPC+). Most interestingly, a cluster of lung progenitor cells (Cluster 7. TM4SF1+/KRT5+/SOX9+) were identified, which will be analyzed in details as below (Fig. 1a and Extended Data Fig. 2).

**Fig. 1.**
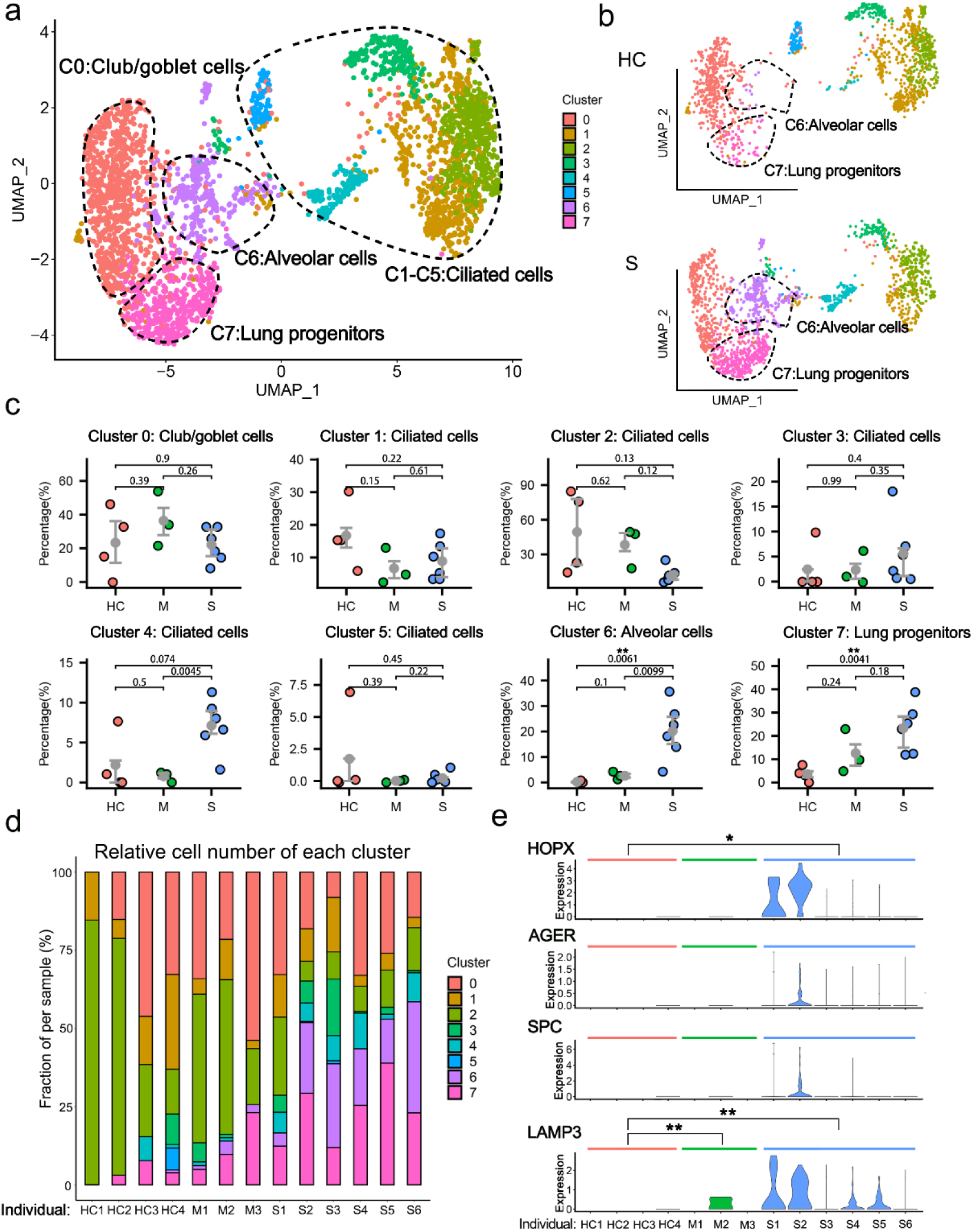
The epithelial cell landscape in the BALF of COVID-19 patients. **a**, The UMAP presentation of the heterogeneous clusters of BALF epithelial cells (all individuals combined, n = 13). **b**, Comparison of UMAP projection of 8 epithelial clusters among healthy controls (HC, n = 4) and severe COVID-19 patients (S, n = 6). **c**, Comparison of the major BALF epithelial cell type proportions in healthy controls (HC), patients with moderate (M) and severe (S) COVID-19 infection. P values were indicated by numbers. **, P<0.01. **d**, The bar plot shows the percentage of epithelial cell clusters in each individual. **e**, The gene expression levels of selected alveolar markers in Cluster 6 from healthy controls (HC, n = 4), moderate cases (M, n = 3) and severe cases (S, n = 6). *P < 0.05, **P < 0.01. P-value adjusted by false discovery rate in MAST.

When we compare the HC group with the other two infected groups, we found significant higher proportion of alveolar cell clusters (Cluster 6) in the BALF of patients with severe infection (Fig. 1b-d). Of note, the HOPX+/AGER+ type I alveolar cells (ATI) and SPC+/LAMP3+ ATII were almost undetectable in the BALFs of healthy control persons due to the tissue integrity of their lungs. In contrast, in the severe COVID-19 patients, both ATI and ATII cell markers were detected in the lavage fluid, probably due to the tissue collapse and desquamation of alveolar cells (Fig. 1e). This phenomenon was not obvious in moderate COVID-19 patients, which was also consistent with previous pathological observation^7^. Therefore, the number of alveolar cells (or the alveolar marker gene expression level) in BALFs could be clinically used to measure the structural integrity of lung, which could serve as an index of disease severity for COVID-19 patients.

In the BALFs of patients with severe infection, we also found significant higher proportions of progenitor cell clusters (Cluster 7) (Fig. 1b-d). Multiple stem/progenitor cell populations have been reported to play critical roles in damage repair after various types of acute lung injury^8^. Among them, a rare population of Wnt-responsive ATII is regarded as the major facultative progenitors^9,10^, which can be specifically marked by TM4SF1 expression in human lung^11^. In current study, we found that in the patients of severe group, the number of TM4SF1+ cells increased remarkably, which implicated the rapid activation of such progenitor cells by tissue damage (Fig. 2a,b). Consistently, we observed co-expression of TM4SF1 and mature alveolar cell markers, AGER (also known as RAGE) and SPC, in a substantial proportion of cells in patients of severe group (Fig. 2c). These results suggested that the TM4SF1+ progenitor cells had the potential to differentiate into mature alveolar cells and regenerate the damaged alveoli of COVID-19 patients.

**Fig. 2.**
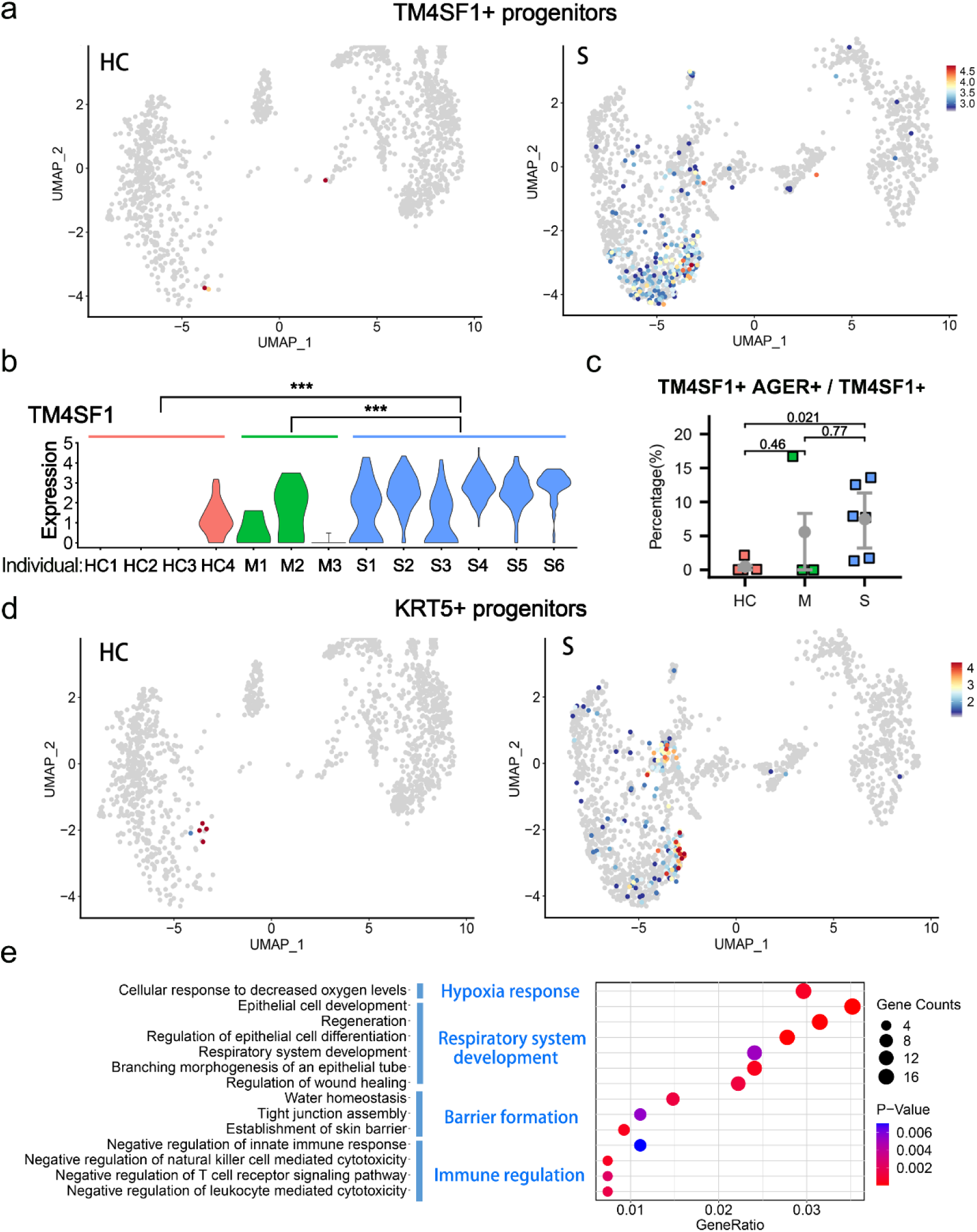
Enrichment of TM4SF1+ and KRT5+ lung progenitors in severe COVID-19 patients. **a**, UMAP plots showing the TM4SF1+ cells in healthy controls (HC, n = 4) and severe COVID-19 patients (S, n = 6). **b**, The gene expression levels of TM4SF1 in Cluster 7 (lung progenitors) from healthy controls (HC), moderate cases (M) and severe cases (S). ***P < 0.001. P-value was adjusted by false discovery rate in MAST. **c**, The various proportions of TM4SF1+ AEGR+ double-positive cells in all TM4SF1+ cells among healthy controls (HC) and patients with moderate (M) and severe (S) COVID-19 infection. P-value was indicated as numbers. **d**, UMAP plots showing the KRT5+ cells in healthy controls (HC, n = 4) and severe COVID-19 patients (S, n = 6). **e**, Gene Ontology enrichment analysis of the differentially expressed genes identified in the KRT5+ cell population.

KRT5+ cells are also reported to have lung stem/progenitor characteristics. Such cells are originated from various primitive progenitors in proximal or distal airways and could expand/migrate to inflamed damaged lung parenchymal region to form “KRT5 pods” once activated by various types of tissue injury including influenza virus infection^12-17^. Recent studies showed that the expanded KRT5 cells could give rise to new pulmonary epithelium, which was now believed to have important epithelial barrier function to protect the lung tissue from further damage^11,17,18^. Specific ablation of the newly expanded KRT5+ cells resulted in persistent hypoxemia, confirming the contribution of these cells in recovery of lung function^15^. In current study, we found that in the patients of severe group, the number of KRT5+ progenitor cells increased remarkably, in together with the elevated expression of another related progenitor marker gene SOX9^19,20^ (Fig. 2d and Extended Data Fig. 2c).

Then we examined the function of genes whose expression level were significantly up-regulated in KRT5+ cells by Gene Ontology analysis. The results showed that KRT5+ cells were responsive to the low oxygen condition of severe COVID-19 patients, and actively participated in the development and generation of respiratory epithelial system. More importantly, such cells highly expressed multiple tissue integrity genes including Claudin1, Claudin4, TJP1, Stratifin, AQP3 and Scnn1A, which were associated with epithelial barrier establishment, tight junction assembly, maintenance of fluid homeostasis and prevention of leukocyte mediated cytotoxicity (Fig. 2e). These functions of KRT5+ cells were rather important for a damaged lung as reconstitution of tight alveolar barrier between atmosphere and fluid-filled tissue is required for recovery of normal gas exchange, while persistent disruption of the alveolar barrier could result in catastrophic consequences including alveolar flooding, “cytokine storm” attack from the circulating leukocytes and subsequent fibrotic scarring^21^.

In order to elucidate the process that how KRT5+ progenitor cells restored mature alveolar barrier in injured lung, we isolated the mouse KRT5+ progenitor cells (previously also named distal airway stem cells)^15^ for transplantation assay as described in Fig. 3a. Briefly, the cell population was trypsinized into single-cell suspension and a single cell-derived pedigree clone was propagated, followed by GFP labeling by lentiviral infection for further analysis. Immunostaining showed that all cultured GFP+ KRT5+ progenitor cells expressed KRT5 and P63 marker genes (Fig. 3b). Then we transplanted the cultured KRT5+ progenitor cells into the bleomycin-injured mouse lung by intratracheal instillation. Transplanted lungs were evaluated at 30 and 90 days afterwards. The result showed substantial persistent engraftment of the GFP+ cells into the host lung, which takes up 2.27% and 2.44% of total lung cells on Day 30 and 90 post transplantation, respectively (Fig. 3c,d). Cryo-sectioning and immunostaining of the host lung demonstrated that some transplanted GFP+ KRT5+ progenitor cells differentiated into alveolar barrier cells (ABCs) in lung parenchyma, establishing a large area of sealed cavities next to consolidated regions, which efficiently prevented CD45+ immune cell infiltration into such area (Fig. 3e and Extended Data Fig. 3a).

**Fig. 3.**
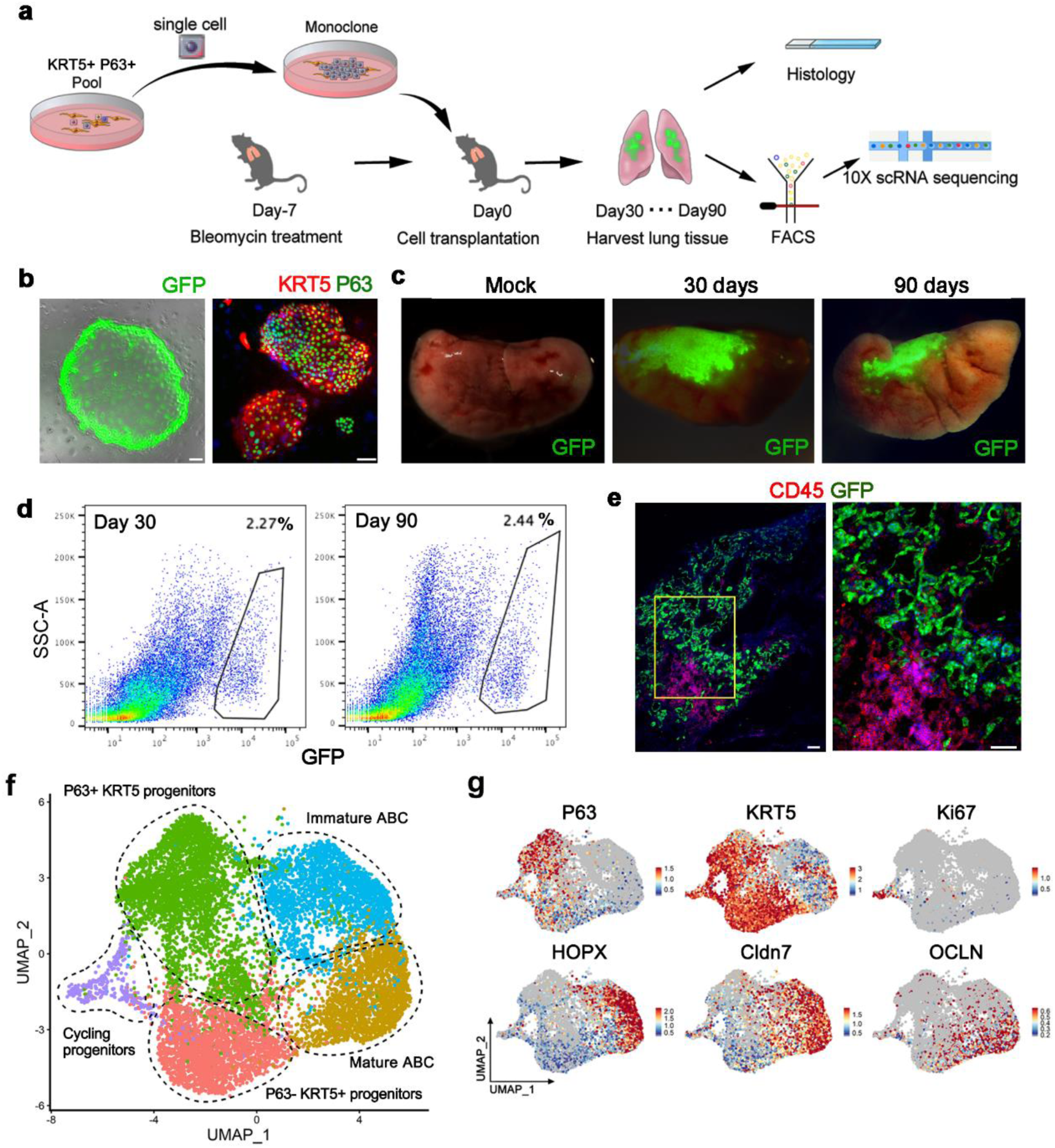
Long-term engraftment of single cell-derived KRT5+ progenitors in host lung. **a**, Schematic showing procedure of single-cell isolation and following analysis. **b**, Cultured cell colonies expressed GFP and progenitor cell markers P63 and KRT5. Scale bar, 100 μm. **c**, Direct GFP imaging of lung lobes following progenitor cell transplantation. **d**, Gating strategy to purify GFP-labeled engrafted cells from transplanted lungs. **e**, Protein immunostaining of CD45 (immune cell) and GFP (engrafted cells) in transplanted lungs at Day 30. Scale bar, 100 μm. **f**, UMAP plot of all engrafted cell clusters 30 and 90 days post transplantation, respectively. ABC, alveolar barrier cell. **g**, UMAP plots showing distinct populations of cells with specific marker expression.

Next we harvested the 30-day and 90-day transplanted lungs and sorted the engrafted GFP+ single cells for scRNA-Seq analysis. Clustering analysis of the sequencing data identified three clusters of KRT5+ progenitor cells, including the most immature P63+ KRT5+ progenitors, the P63-KRT5+ progenitors and the actively cycling Ki67+ KRT5+ progenitors. We also identified two KRT5-differentiated ABCs which both highly expressed early alveolar marker HOPX and paracellular adhesion marker Claudin7. One of the two ABC clusters highly expressing tight junction marker Occludin (OCLN) was supposed to be more mature than the other one (Fig. 3f,g). Gene ontology analysis demonstrated that the P63+ KRT5+ progenitors highly expressed multiple genes whose function was related to cell stemness maintenance, chemotaxis and inhibition of cell apoptosis. In contrast, the mature ABC highly expressed multiple genes (including HOPX, SPD, OCLN, CDH1, Claudin7, Claudin4, AQP3, etc.) whose function was related to lung morphogenesis, tight junction assembly and water homeostasis (Extended Data Fig. 3b,c).

To further dissect the lineage relationship between different clusters of engrafted cells, we performed the Monocle pseudo-time analysis based on the scRNA-Seq data. The result indicated that the P63+ KRT5+ progenitors could differentiate into P63-KRT5+ progenitors and immature ABCs, which eventually give rise to OCLN+ mature ABCs (Fig. 4a,b). In consistent with the pseudo-time analytical data, we noticed that comparing to the 30-day engrafted cells, the 90-day engrafted cells had relatively more mature ABCs, less immature ABCs and less P63-KRT5+ progenitors (Fig. 4c). Altogether such data revealed that the tight alveolar barrier would be gradually established by KRT5+ progenitor cell differentiation.

**Fig. 4.**
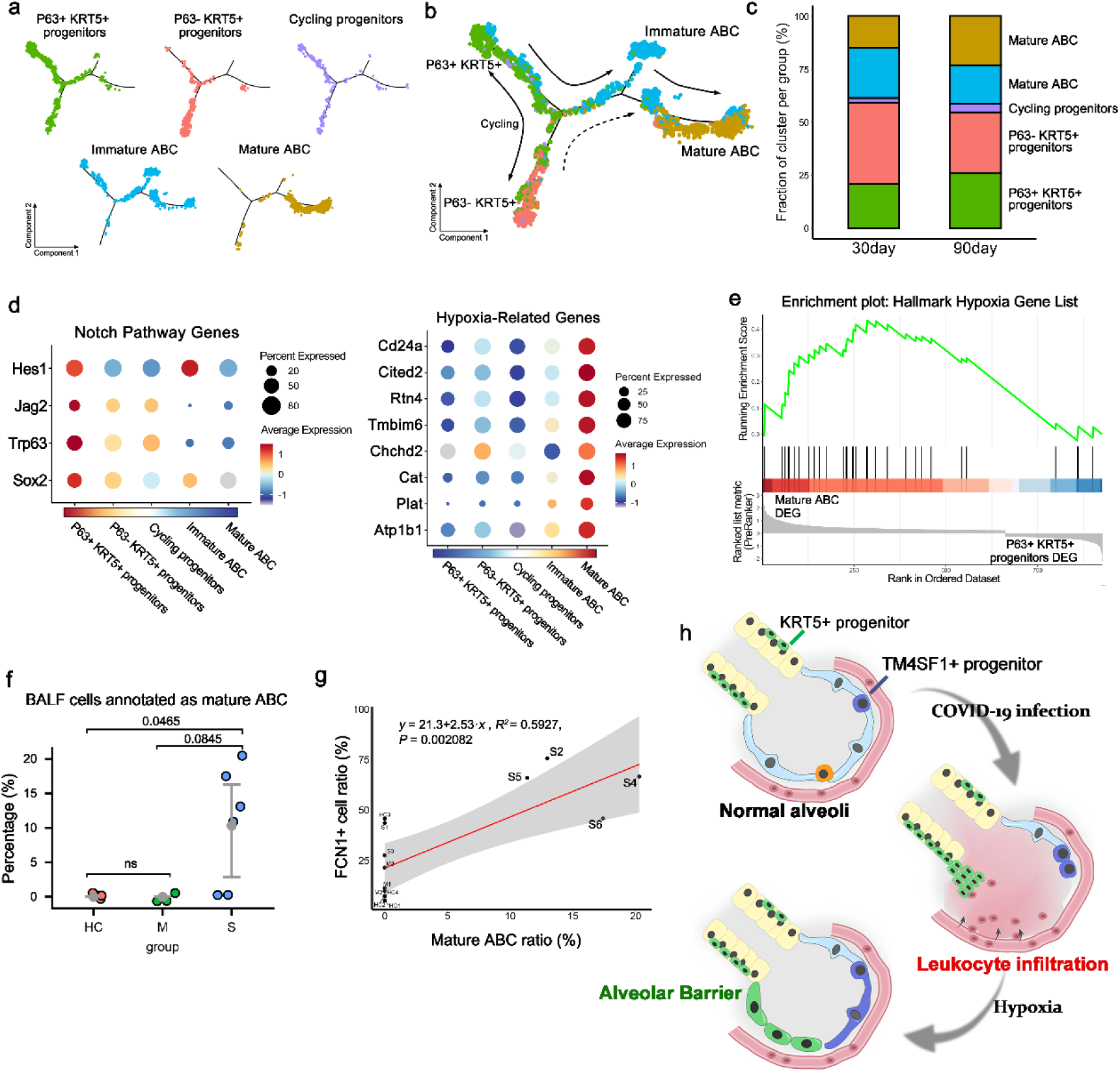
KRT5+ progenitors give rise to alveolar barrier cells. **a**, The pseudotime trajectory showing the distribution of distinct cell clusters. **b**, Pseudotime trajectory analysis shows the putative differentiation paths from P63+ KRT5+ progenitors to mature alveolar barrier cells. **c**, The bar plot showing the percentages of distinct cell clusters in 30 day and 90 day sample. **d**, Expression of selected Notch pathway and hypoxia related-genes is shown in different cell types. **e**, Gene Set Enrichment Analysis is performed using the Hallmark Hypoxia Gene Set with differentially expressed genes between P63+ KRT5+ progenitors and mature ABC. **f**, The percentage of BALF cells annotated as mature alveolar barrier cells among healthy controls (HC) and patients with moderate(M) and severe (S) COVID-19 infection. **g**, Correlation analysis between percentage of FCN1+ cells in all macrophages and percentage of BALF cells annotated to mature alveolar barrier cells in each sample. Each dot corresponds to each sample (n=13). **h**, Schematic of alveolar barrier establishment post COVID-19 infection.

Next we asked which molecular signaling pathways were involved in the establishment of alveolar barrier. Previous studies indicated that Notch signaling is critical for activation of P63+ KRT5+ progenitors in lung, but persistent Notch signaling prevents further differentiation of cells ^16^. Consistently, here we found that the expression of multiple Notch pathway component genes was up-regulated in P63+ KRT5+ progenitors but gradually down-regulated when the cells were differentiating to mature ABC (Fig. 4d). In addition, sense of low oxygen level through hypoxia pathway is known to be critical for the expansion of KRT5+ progenitors^22^. Here we found that the hypoxia pathway component gene expressions were relatively low in P63+ KRT5+ progenitors but were gradually up-regulated when the cells were differentiating to ABC (Fig. 4d). Further gene set enrichment analysis (GSEA) also demonstrated that the expression of the hallmark hypoxia gene list in Molecular Signatures Database (MSigDB) was significantly enriched in mature ABC (P=0.018) (Fig. 4e). Altogether these data showed that Notch and hypoxia pathways were important for the regulation of progenitor cell fate.

Lastly, we mapped our scRNA-Seq data of transplanted progenitor cells back to the COVID-19 BALF scRNA-Seq data to investigate whether any mature alveolar barrier has been established in patients. We extracted the mature ABC gene expression signature and evaluate the signature signal in COVID-19 patients. The result showed that approximately 10% of the epithelial cells in BALF of severe patients could be annotated as mature ABC, and such ABC only existed in 4 severe patients (S2, S4, S5 and S6) but not in healthy persons or other patients (Fig. 4f). So why the 4 patients have much more mature ABC than others? Interestingly, we found that generally the individuals’ mature ABC cell numbers were positively correlated with their FCN1+ macrophage cell numbers in BALF (P=0.002), and the patient S2, S4, S5 and S6 had much more FCN1+ macrophages in BALF than most of the other individuals (Fig. 4g). FCN1+ macrophages were reported to be highly pro-inflammatory and responsible for the tissue damage in COVID-19 patients^5^. Therefore, it seems that the establishment of new alveolar barrier was closely associated with severity of tissue damage and inflammation at individual level.

Altogether, our current studies uncovered a possible mechanism of lung repair after severe SARS-CoV-2 infection. In those severe patients, the virus infection led to alveolar cell death and leakage of epithelial barrier, which resulted in massive flood of proteins-enriched fluids and leukocytes into alveolar cavity, which finally leading to pulmonary edema and tissue hypoxia. In this situation, as repair backups, KRT5+ and TM4SF1+ progenitor cell could be activated by the hypoxia and other multiple microenvironment signals to expand, migrate and differentiate into new functional cells. Eventually, the KRT5+ progenitor-derived ABCs could rapidly restore new epithelial barriers to cover the denuded alveoli and sealed the leakage. Simultaneously, the TM4SF1+ progenitor cells could gradually regenerate new functional ATII and ATI cells. The synergic act of two types of progenitor cells could make timely repair of alveoli in an acute injury scenario (summarized in Fig. 4h). Of course, there could be more resident or circulating stem/progenitor cells working in concert with them to achieve maximal repair. Of note, our current studies were based on limited sample data. Future study within a larger patient cohort, could further facilitate our understanding of the repair process of COVID-19 patients, and development of potential progenitor cell-based therapeutic strategies.

## Methods

### scRNA-seq analysis of BALF cells

Public datasets (GEO: GSE145926) ^5^and (GEO: GSE128033) ^6^which contain scRNA-seq data from BALF cells from three patients with moderate COVID-19 (M1-M3), six patients with severe/critical COVID-19 (S1-S6), three healthy controls (HC1-HC3) and one fresh BALF (GSM3660650) from a lung transplant donor (HC4) samples were used for bioinformatic analysis. Seurat v.3 was used for quality control. The following criteria were then applied to each cell of all nine patients and four healthy controls: gene number between 200 and 6,000, UMI count > 1,000 and mitochondrial gene percentage < 0.1. After filtering, a total of 66,452 cells were left for the following analysis. A filtered gene-barcode matrix of all samples was integrated with Seurat v.3 to remove batch effects across different donors. In parameter settings, the first 50 dimensions of canonical correlation analysis (CCA) and principal-component analysis (PCA) were used.

The filtered gene-barcode matrix was first normalized using ‘LogNormalize’ methods in Seurat v.3 with default parameters. The top 2,000 variable genes were then identified using the ‘vst’ method in Seurat FindVariableFeatures function. Variables ‘nCount_RNA’ and ‘percent.mito’ were regressed out in the scaling step and PCA was performed using the top 2,000 variable genes. Then UMAP was performed on the top 50 principal components for visualizing the cells. Meanwhile, graph-based clustering was performed on the PCA-reduced data for clustering analysis with Seurat v.3. The resolution was set to 1.2 to obtain a finer result.

Epithelial cells were re-clustered with Seurat v.3. Epithelial cells of all samples were re-clustered using the same parameter mentioned above in the clustering step and parameter resolution was set to 0.3. Immune cell clusters were removed and the other cells were re-clustered using the same parameter mentioned above in the clustering step and parameter resolution was set to 0.4. MAST^23^ in Seurat v.3 (FindAllMarkers function) was used to perform differential gene expression analysis. For each cluster of epithelial cells, DEGs were generated relative to all of the other cells. A gene was considered significant with adjusted P < 0.05 (P values were adjusted by false discovery rate in MAST).

For analysis of engrafted GFP+ cells, single cells were captured and barcoded in 10x Chromium Controller (10x Genomics). Subsequently, RNA from the barcoded cells was reverse-transcribed and sequencing libraries were prepared using Chromium Single Cell 3’v3 Reagent Kit (10x Genomics) according to the manufacturer’s instructions. Sequencing libraries were loaded on an Illumina NovaSeq with 2×150 paired-end kits at Novogene, China. Raw sequencing reads were processed using the Cell Ranger v.3.1.0 pipeline from 10X Genomics. In brief, reads were demultiplexed, aligned to the human GRCh38 genome and UMI counts were quantified per gene per cell to generate a gene-barcode matrix. Data were aggregated and normalized to the same sequencing depth, resulting in a combined gene-barcode matrix of all samples.

Post-processing, including filtering by number of genes and mitochondrial gene content expressed per cell, was performed using the Seurat v.3. Genes were filtered out that were detected in less than 3 cells. A global-scaling normalization method ‘LogNormalize’ was used to normalize the data by a scale factor (10,000). Next, a subset of highly variable genes was calculated for downstream analysis and a linear transformation (ScaleData) was applied as a pre-processing step. Principal component analysis (PCA) dimensionality reduction was performed with the highly variable genes as input in Seurat function RunPCA. The top 20 significant PCs were selected for two-dimensional t-distributed stochastic neighbor embedding (tSNE), implemented by the Seurat software with the default parameters. FindCluster in Seurat was used to identify cell clusters.

### Gene functional annotation

Gene Ontology (GO) enrichment analysis and Gene Set Enrichment Analysis (GSEA) of differentially expressed genes was implemented by the ClusterProfiler R package ^24^. GO terms with corrected P value less than 0.05 were considered significantly enriched by differentially expressed genes. Dot plots were used to visualize enriched terms by the enrichplot R package. For hypoxia gene analysis, the hallmark gene sets in MsigDB ^25^ were used for annotation.

### Single cell trajectory analysis

To construct single cell pseudotime trajectory and to identify genes that change as the cells undergo transition, Monocle2 (version 2.4.0) algorithm ^26^ was applied to our datasets. Genes for ordering cells were selected if they were expressed in ≥ 1% cells, their mean expression value was ≥ 0.3 and dispersion empirical value was ≥ 1. Based on the ‘DDRTree ‘method, the data was reduced to two dimensional, and then the cells were ordered along the trajectory.

### Correlation analysis of BALF cells and engrafted cells

The scHCL Model^27^ ^2726^was used to assess the similarity between BALF cells and engrafted cells. The expression patterns of highly variable genes of each cluster of engrafted cells (eg. mature ABC) was taken as reference signatures to annotate BALF cells. Pearson correlation score was calculated between each cell type of scRNA-seq data from engrafted cells and each cell of scRNA-seq data from BALF cells. Final annotation of each BALF cell was based on the highest correlation score. Linear regression analysis was performed between ratio of FCN1+ macrophage cell numbers in BALF and ratio of the epithelial cells in BALF annotated as mature ABC with ‘lm’ method.

### Cell culture

Mouse lung P63+KRT5+ progenitor cells were isolated and cultured as previously described ^15^. Briefly, lung lobes were collected and processed into a single-cell suspension by protease, trypsin and DNaseI. Dissociated cells were passed through 70-μm nylon mesh, washed twice with cold F12 medium and then cultured onto feeder cells(irradiated 3T3-J2 feeder cells) in culture medium including DMEM/F12, 10% FBS (Hyclone, Australia), Pen/Strep, amphotericin, and growth factor cocktail as previously described. Cells were grown in a humidified atmosphere of 7.5% (v/v) CO2 at 37 °C. To generate monoclonal cell, cells were processed and diluted into single-cell suspension. One single cell was aspirated and isolated by pipette under microscopy, and then transferred into 96 well plate to expand.

### Animal experiments

C57/B6 mice (6–8 weeks) were purchased from Shanghai SLAC Laboratory Animal Co., Ltd. (China) and housed in specific pathogen-free conditions within an animal care facility (Center of Laboratory Animal, Tongji University, Shanghai, China). All animals were cared for in accordance with NIH guidelines, and all animal experiments were performed under the guidance of the Institutional Animal Care and Use Committee of Tongji University. For the lung injury mouse model, bleomycin was intratracheally administrated to isoflurane-anesthetized mice at a concentration of 3U/kg seven days prior transplantation. GFP labeled cells suspended in 40uL DMEM (1 million cells per mouse) were intratracheally transplanted. At distinct time points, mice were sacrificed and their lung tissue were harvested to detect GFP signal by fluorescence stereomicroscope (MVX10, Olympus, Japan).

### Histology

Tissues were fixed with 4% PFA for 2 hours at room temperature and at 4°C overnight, followed by embedding in OCT. Immunohistochemistry was performed following heat antigen retrieval methods and stained with the following antibodies. GFP (goat, Abcam, 1:1000), Krt5 (rabbit, Thermo Fisher, 1:200), P63 (mouse, abcam, 1:100) and CD45 (rabbit, abcam, 1:200). Primary antibody was incubated at 4°C overnight, and second antibody at room temperature for 2h.

### FACS sorting of engrafted GFP+ single cells

Transplanted lung was collected and immersed in cold F12 medium with 5% FBS, followed by being minced into small pieces and digested with dissociation buffer (F12, 1mg/ml protease, 0.005% trypsin and 10ng/ml DNase I) on shaker in 37 degree for 1hr. Dissociated cells was filtered through 100-μm cell strainer, and Red Blood Cell Lysis Buffer was used to remove erythrocyte. Cell pellets were resuspended in DMEM containing with 1% FBS following washing twice, and then passed over 30-μm strainers. Sorting and subsequent quantification were performed on BD FACS Arial cytometers. GFP+ cells were gated using SSC-A vs FSC-A, FSC-H vs FSC-W, and SSC-H vs SSC-W gates, followed by SSC-A vs FITC-A gate.

### Statistics

Differences of median percentage between healthy controls, moderate and severe groups of all cell types, TM4SF1+ AEGR+ cells in all TM4SF1+ cells and BALF cells annotated to mature ABCs were compared using a Student’s t-test (two-sided, unadjusted for multiple comparisons) with R ggpubr v.0.2.5. Differences of gene expression levels between healthy controls, moderate and severe groups were compared using MAST in Seurat v.3. A gene was considered significant with adjusted P < 0.05 (P values were adjusted by false discovery rate in MAST).

## Acknowledgements

W. Zuo and T. Zhang designed the project; Z. Zhao, Y. Zhao, Y. Zhou, and X. Wang performed the analysis; Z. Zhao, Y. Zhao, Y. Zhou and W. Zuo drafted the manuscript. We thank Prof. Zheng Zhang group for sharing their original COVID-19 BALF scRNA-Seq dataset to public. This work was funded by the National Key Research and Development Program of China (2017YFA0104600 to W. Zuo), National Science Foundation of China (81770073 to W. Zuo), Shanghai Science and Technology Talents Program (19QB1403100 to W.Zuo), Tongji University (Basic Scientific Research Interdisciplinary Fund and 985 Grant to W. Zuo) and Guangzhou Medical University annual grant to W. Zuo. We salute to medical workers who sacrificed their lives in fight against the COVID-19 pandemic.

---

**Extended Data Fig. 1.**
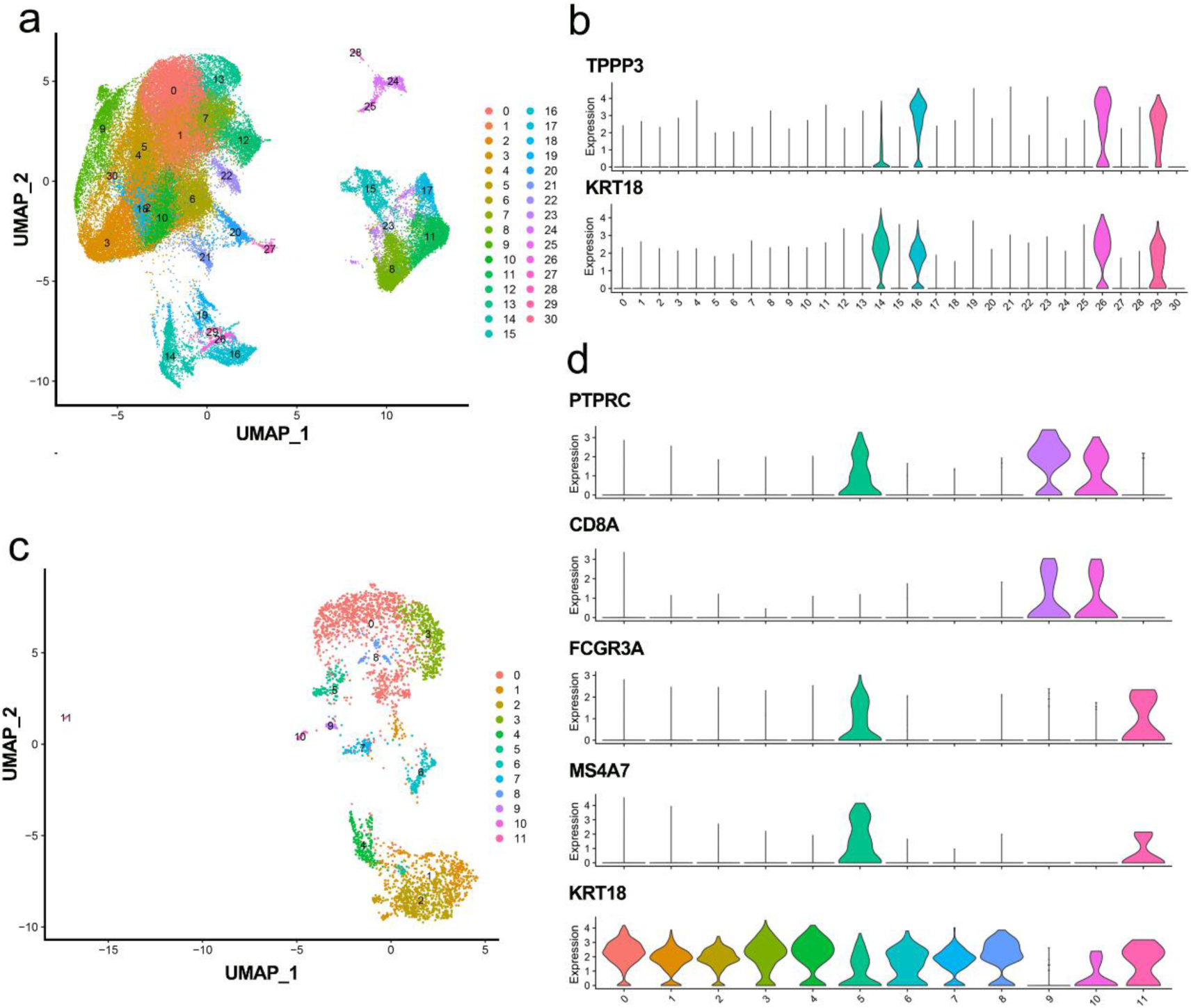
scRNA-seq analysis of all BALF cells. **a**, UMAP presentation of all major cell types in BALF (n = 13). **b**, The gene expression levels of epithelial cell markers in each cluster (n = 13). **c**, The original UMAP presentation of major cell types in BALF epithelial cells without removing leukocyte-engulfed cells. (n = 13). **d**, The gene expression levels of immune cell markers in original BALF epithelial cells. Cluster 5 (FCGR3A+ MS4A7+), 9 (PTPRC+ CD8A+), 10 (PTPRC+ CD8A+) and 11 (FCGR3A+ MS4A7+) were excluded in further analysis (n = 13).

**Extended Data Fig. 2.**
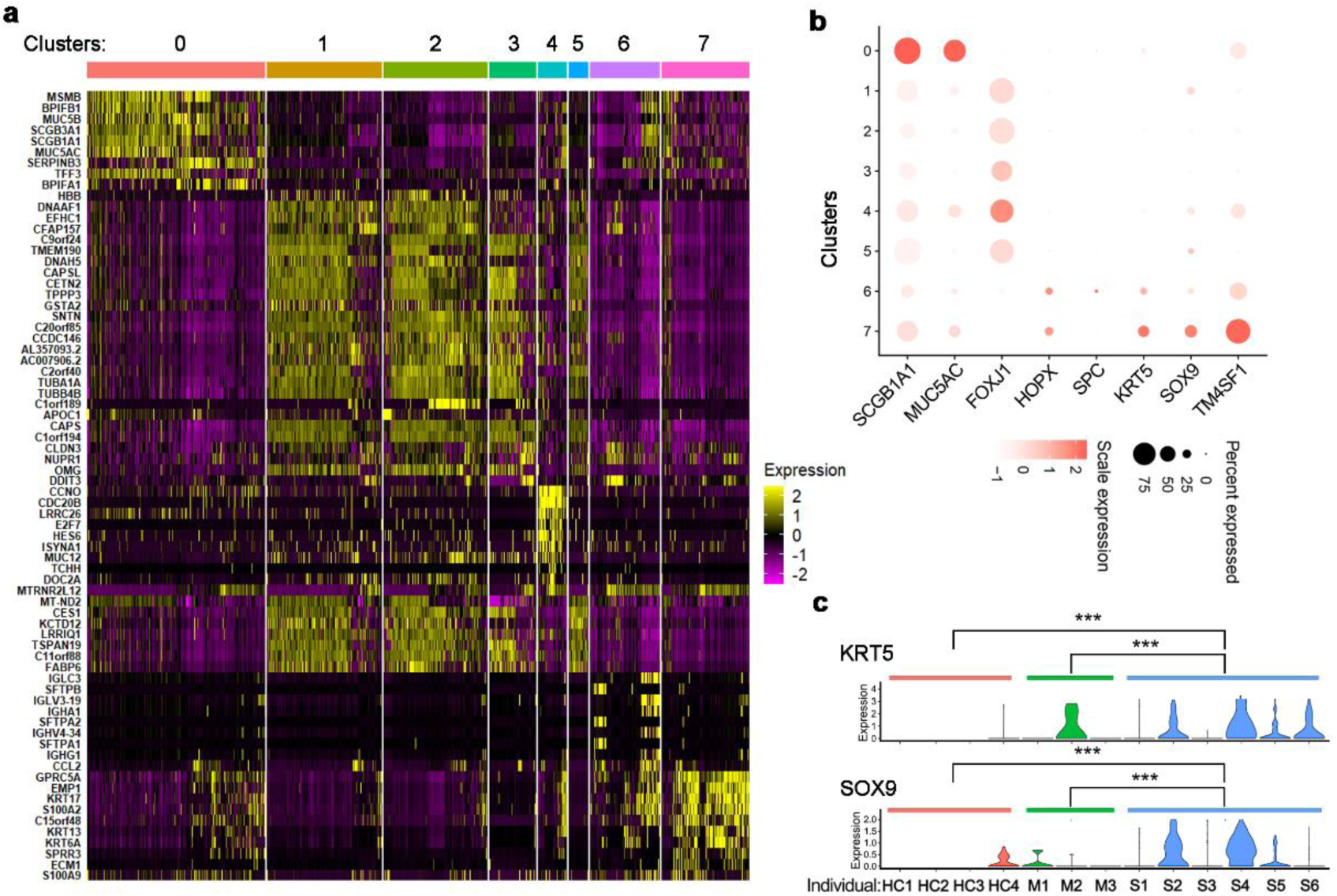
Identification of clusters from BALF epithelial cells. **a**, Heat map of the top ranked genes highly expressed in each cluster. Color scheme is based on z-score distribution from –2 (purple) to 2 (yellow). **b**, A dot plot showing expression of hallmark genes by different cell types in BALF epithelial cells. (n = 13). **c**, The gene expression levels of KRT5 and SOX9 in cluster 7 lung progenitors from healthy controls (n = 4), moderate cases (n = 3) and severe cases (n = 6). ***P < 0.001.

**Extended Data Fig. 3.**
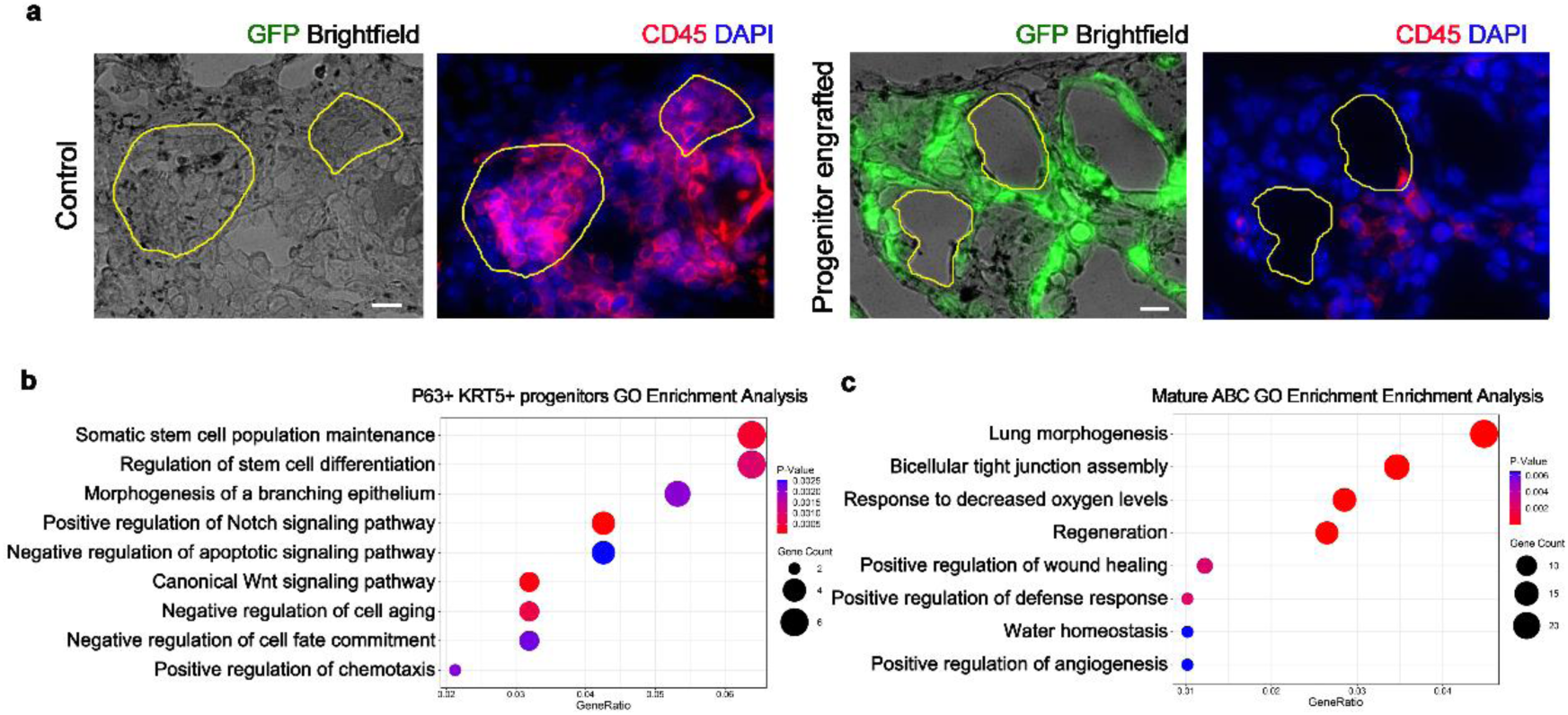
Transplanted KRT5+ progenitors differentiate into alveolar barrier cells. **a**, Massive immune cell infiltration into alveolar cavity in non-transplanted control lung but not in transplanted lung with GFP+ progenitor engrafted. Scale bar, 25 μm. **b and c**, Gene Ontology enrichment analysis of the differentially expressed genes identified in P63+ KRT5+ progenitors and mature alveolar barrier cells.

## References

1 Mo, X. et al. Abnormal pulmonary function in COVID-19 patients at time of hospital discharge. The European respiratory journal 55, doi: 10.1183/13993003.01217-2020 (2020).

2 Tian, S. et al. Pathological study of the 2019 novel coronavirus disease (COVID-19) through postmortem core biopsies. Modern pathology : an official journal of the United States and Canadian Academy of Pathology, Inc 33, 1007–1014, doi: 10.1038/s41379-020-0536-x (2020).

3 Wu, J. H. et al. [Pathological changes of fatal coronavirus disease 2019 (COVID-19) in the lungs: report of 10 cases by postmortem needle autopsy]. Zhonghua bing li xue za zhi = Chinese journal of pathology 49, 568–575, doi: 10.3760/cma.j.cn112151-20200405-00291 (2020).

4 Xu, Z. et al. Pathological findings of COVID-19 associated with acute respiratory distress syndrome. The Lancet. Respiratory medicine 8, 420–422, doi: 10.1016/S2213-2600(20)30076-X (2020).

5 Liao, M. et al. Single-cell landscape of bronchoalveolar immune cells in patients with COVID-19. Nat Med 26, 842–844, doi: 10.1038/s41591-020-0901-9 (2020).

6 Morse, C. et al. Proliferating SPP1/MERTK-expressing macrophages in idiopathic pulmonary fibrosis. The European respiratory journal 54, doi: 10.1183/13993003.02441-2018 (2019).

7 Tian, S. et al. Pulmonary Pathology of Early-Phase 2019 Novel Coronavirus (COVID-19) Pneumonia in Two Patients With Lung Cancer. Journal of thoracic oncology : official publication of the International Association for the Study of Lung Cancer 15, 700–704, doi: 10.1016/j.jtho.2020.02.010 (2020).

8 Basil, M. C. et al. The Cellular and Physiological Basis for Lung Repair and Regeneration: Past, Present, and Future. Cell Stem Cell 26, 482–502, doi: 10.1016/j.stem.2020.03.009 (2020).

9 Barkauskas, C. E. et al. Type 2 alveolar cells are stem cells in adult lung. Journal of Clinical Investigation 123, 3025–3036 (2013).

10 Nabhan, A. N., Brownfield, D. G., Harbury, P. B., Krasnow, M. A. & Desai, T. J. Single-cell Wnt signaling niches maintain stemness of alveolar type 2 cells. Science 359, 1118–1123, doi: 10.1126/science.aam6603 (2018).

11 Zacharias, W. J. et al. Regeneration of the lung alveolus by an evolutionarily conserved epithelial progenitor. Nature 555, 251–255, doi: 10.1038/nature25786 (2018).

12 Yang, Y. et al. Spatial-Temporal Lineage Restrictions of Embryonic p63(+) Progenitors Establish Distinct Stem Cell Pools in Adult Airways. Dev Cell 44, 752–761 e754, doi: 10.1016/j.devcel.2018.03.001 (2018).

13 Ray, S. et al. Rare SOX2(+) Airway Progenitor Cells Generate KRT5(+) Cells that Repopulate Damaged Alveolar Parenchyma following Influenza Virus Infection. Stem cell reports 7, 817–825, doi: 10.1016/j.stemcr.2016.09.010 (2016).

14 Kumar, P. A. et al. Distal airway stem cells yield alveoli in vitro and during lung regeneration following H1N1 influenza infection. Cell 147, 525–538 (2011).

15 Zuo, W. et al. p63+Krt5+ distal airway stem cells are essential for lung regeneration. Nature 517, 616–620 (2015).

16 Vaughan, A. E. et al. Lineage-negative progenitors mobilize to regenerate lung epithelium after major injury. Nature 517, 621–625 (2015).

17 Kathiriya, J. J., Brumwell, A. N., Jackson, J. R., Tang, X. & Chapman, H. A. Distinct Airway Epithelial Stem Cells Hide among Club Cells but Mobilize to Promote Alveolar Regeneration. Cell Stem Cell 26, 346–358 e344, doi: 10.1016/j.stem.2019.12.014 (2020).

18 Kanegai, C. M. et al. Persistent Pathology in Influenza-Infected Mouse Lungs. American journal of respiratory cell and molecular biology 55, 613–615, doi: 10.1165/rcmb.2015-0387LE (2016).

19 Ma, Q. et al. Regeneration of functional alveoli by adult human SOX9 + airway basal cell transplantation. Protein & Cell 9, 267–282 (2018).

20 Nichane, M. et al. Isolation and 3D expansion of multipotent Sox9(+) mouse lung progenitors. Nature methods 14, 1205–1212, doi: 10.1038/nmeth.4498 (2017).

21 Yanagi, S., Tsubouchi, H., Miura, A., Matsumoto, N. & Nakazato, M. Breakdown of Epithelial Barrier Integrity and Overdrive Activation of Alveolar Epithelial Cells in the Pathogenesis of Acute Respiratory Distress Syndrome and Lung Fibrosis. BioMed research international 2015, 573210, doi: 10.1155/2015/573210 (2015).

22 Xi, Y. et al. Local lung hypoxia determines epithelial fate decisions during alveolar regeneration. Nature cell biology 19, 904–914, doi: 10.1038/ncb3580 (2017).

23 Finak, G. et al. MAST: a flexible statistical framework for assessing transcriptional changes and characterizing heterogeneity in single-cell RNA sequencing data. Genome biology 16, 278, doi: 10.1186/s13059-015-0844-5 (2015).

24 Yu, G., Wang, L. G., Han, Y. & He, Q. Y. clusterProfiler: an R package for comparing biological themes among gene clusters. Omics : a journal of integrative biology 16, 284–287, doi: 10.1089/omi.2011.0118 (2012).

25 Liberzon, A. et al. The Molecular Signatures Database (MSigDB) hallmark gene set collection. Cell systems 1, 417–425, doi: 10.1016/j.cels.2015.12.004 (2015).

26 Trapnell, C. et al. The dynamics and regulators of cell fate decisions are revealed by pseudotemporal ordering of single cells. Nature biotechnology 32, 381–386, doi: 10.1038/nbt.2859 (2014).

27 Han, X. et al. Construction of a human cell landscape at single-cell level. Nature 581, 303–309, doi: 10.1038/s41586-020-2157-4 (2020).

